# Chronic infections can generate SARS-CoV-2-like bursts of viral evolution without epistasis

**DOI:** 10.1101/2024.10.06.616878

**Authors:** Edwin Rodríguez-Horta, John Strahan, Aaron R. Dinner, John P. Barton

**Affiliations:** Department of Computational and Systems Biology, University of Pittsburgh School of Medicine, USA; Group of Complex Systems and Statistical Physics, Department of Theoretical Physics, Physics Faculty, University of Havana, Cuba; Department of Chemistry and James Franck Institute, University of Chicago, Chicago, Illinois 60637, USA

## Abstract

Multiple SARS-CoV-2 variants have arisen during the first years of the pandemic, often bearing many new mutations. Several explanations have been offered for the surprisingly sudden emergence of multiple mutations that enhance viral fitness, including cryptic transmission, spillover from animal reservoirs, epistasis between mutations, and chronic infections. Here, we simulated pathogen evolution combining within-host replication and between-host transmission. We found that, under certain conditions, chronic infections can lead to SARS-CoV-2-like bursts of mutations even without epistasis. Chronic infections can also increase the global evolutionary rate of a pathogen even in the absence of clear mutational bursts. Overall, our study supports chronic infections as a plausible origin for highly mutated SARS-CoV-2 variants. More generally, we also describe how chronic infections can influence pathogen evolution under different scenarios.

## Introduction

During the SARS-CoV-2 pandemic, multiple variants of concern (VOC) have arisen and spread widely throughout the human population, driving waves of infections and mortality ^1–4^. The spread of new VOCs has been facilitated by their ability to evade adaptive immunity developed by previous infections or vaccines ^5,6^. VOC mutations can also increase virus transmissibility in other ways, such as by improving the receptor binding ability of the viral Spike protein or increasing viral load ^5,6^.

A singular and unexpected characteristic of early VOCs has been their abrupt emergence. New variants such as Alpha, Delta, and Omicron appeared bearing many mutations that had not been previously observed, seemingly making a large evolutionary leap compared to co-circulating variants. This phenomenon is surprising given the tight transmission bottlenecks inferred for SARS-CoV-2 (refs. ^7,8^). During acute infections, few mutations are produced and even fewer are expected to be passed on in new infections ^7,8^. In principle, one would then expect viral evolution to proceed through the gradual accumulation of advantageous (i.e., transmission-increasing) mutations.

Multiple hypotheses have been put forward to explain the sudden appearance of a new, highly transmissible variant with a large number of novel mutations ^9^. One possibility is cryptic transmission, where undetected circulation in humans allows for long-term viral evolution ^10–12^. However, given the 1–12 number of novel mutations observed in VOCs, this scenario would require that variants remain undetected for long periods of time. Circulation in animal reservoirs, followed by subsequent spillover to humans, could also explain the sudden appearance of VOCs with many mutations ^13–16^. As an alternative to hidden circulation in unobserved human or animal populations, epistasis (i.e., non-additive effects of mutations on viral transmissibility) has been cited as a possible factor underlying VOC emergence ^17–20^. If multiple mutations are needed to confer a significant fitness advantage to the virus, then mutants with a small number of mutations may not be observed at high frequencies in humans.

Based on clinical data, chronic SARS-CoV-2 infections have emerged as a plausible source of highly divergent variants. Typical, acute SARS-CoV-2 infections resolve within days to weeks. However, in some individuals, chronic infections can persist for months. Chronically infected individuals often have compromised immune systems that are unable to fully clear infections ^21–24^. During chronic infections, there is sufficient time for SARS-CoV-2 to generate multiple mutations, which can rise in frequency and ultimately fix in the viral population within that individual. Genomic analyses have shown that the rate of accumulation of mutations within chronically infected individuals is higher than the rate of SARS-CoV-2 evolution between individuals ^25^. In addition, VOC mutations have been observed in chronically-infected individuals ^25,26^. Accelerated selection of antibody evasion mutations has also been observed in long-term infections treated with monoclonal antibodies or convalescent plasma ^27^.

Given the potential importance of chronic infections in the evolution of SARS-CoV-2, mathematical modelers have begun exploring its anticipated epidemiological effects in theory and simulations ^17,18,28,29^. Recent work has incorporated immunocompromised hosts into susceptible/infected/recovered (SIR) epidemiological models that also include some component of within-host viral evolution ^17,28^. Smith and Ashby predicted that large jumps in the proportion of novel variants should only be observed when there is a significant amount of epistasis between immune escape mutations and a sufficient proportion of the population is immunocompromised ^17^. In other words, the role of immunocompromised hosts in this model is to allow the virus to cross epistatic fitness valleys. Additional work has also considered fitness valley crossing for infections of different durations, but without modeling effects on transmission ^29^.

In an extensive study, Ghafari et al. considered the effects of chronic infection on the emergence of highly transmissible VOCs ^18^. In their model, VOC mutations fix at a constant rate within chronically infected hosts. They consider multiple fitness landscapes for transmission between individuals, including models where VOC mutations make equal, additive contributions to transmission and “plateau-crossing” models where individual mutations have small effects until a critical number are accumulated. They concluded that chronic infection could facilitate the emergence of VOCs, defined as variants with specific transmission-increasing mutations, especially with plateau-crossing fitness landscapes.

Here, we develop a generic model of pathogen evolution, coupling evolution within hosts and transmission between individuals. The primary goal of our model is to understand how chronic infections can affect pathogen evolutionary dynamics over long times. Using transmission effects of mutations inferred from SARS-CoV-2 data ^30^, we show that bursts of mutations like those observed during the pandemic can occur even with a simple, additive fitness landscape. In particular, we explore how the within-host mutation rate, typical duration of infection, and fraction of infections that are chronic affect the likelihood of mutational bursts. We find that bursty evolution is especially likely when the acute infection time is short compared to the duration of chronic infections. Our results highlight scenarios in which chronic infections produce evolutionary dynamics that are qualitatively different from those that are observed in most simple evolutionary models.

## Results

### Model of pathogen evolution within and between hosts

The global evolutionary dynamics of pathogens such as SARS-CoV-2 are a consequence of processes that occur within and between infected individuals. Evolution within individuals generates a genetically diverse cloud or “quasispecies” of variants ^31–33^. Differential transmission of variants between hosts ultimately results in pathogen evolution across individuals. We include both levels of evolution in our model (**Fig. 1**).

**Fig. 1.**
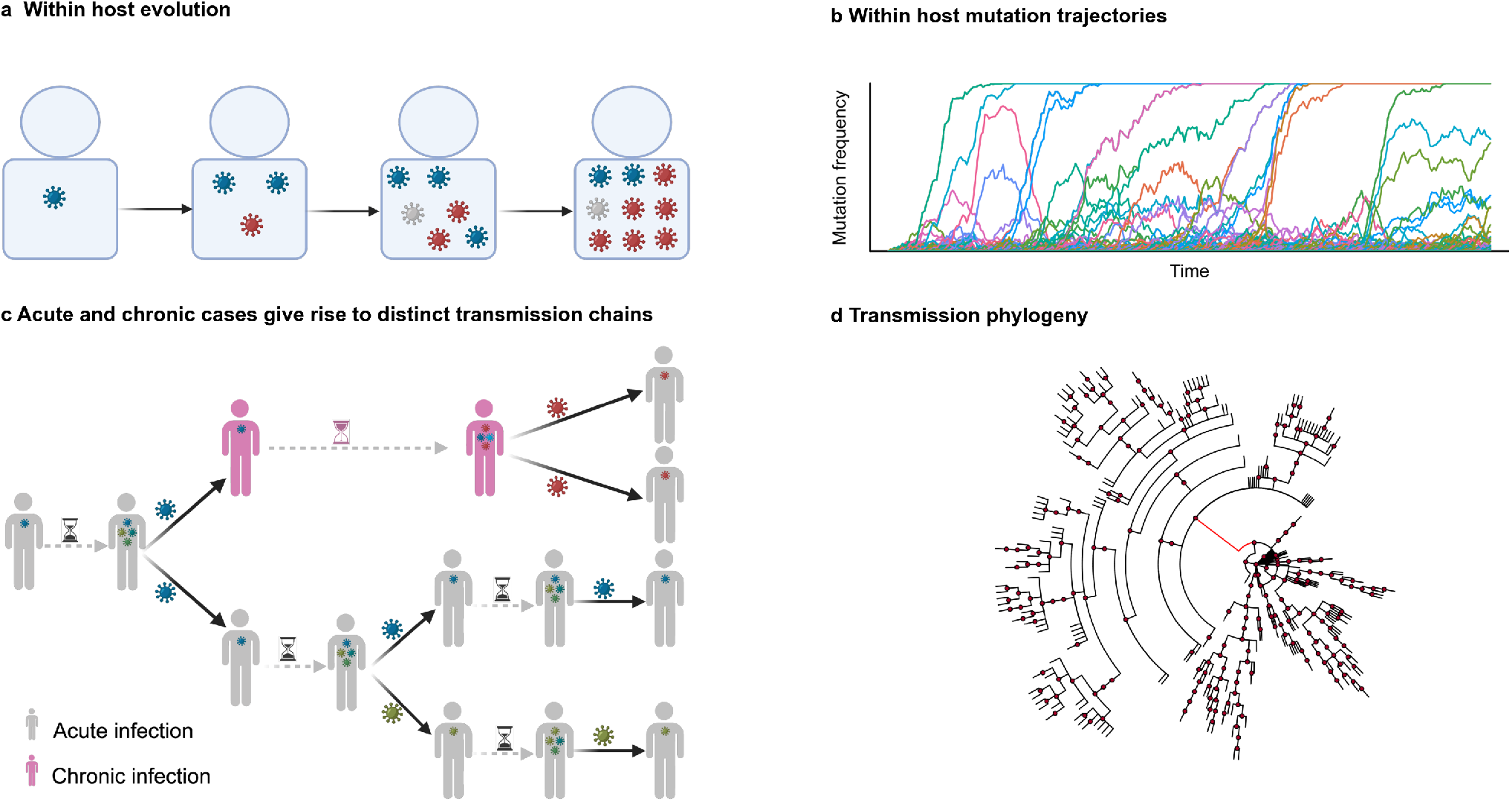
Global evolutionary model for intra-host and between-host levels of evolution. **a**, The viral population, begins with a single starting genotype and undergoes discrete and non-overlapping generations of Wright-Fisher evolution subject to mutation, selection, and genetic drift. **b**, Example frequency of mutations over time during intrahost evolution. **c**, The population of individuals comprises patients with acute infection and patients with chronic infections. Viral diversity is generated during intrahost evolution. During transmission between acutely infected hosts, most mutations are lost due to tight transmission bottlenecks. Chronic infection can allow for the evolution and transmission of a highly divergent variant. **d**, Example phylogeny for between-host transmission, including transmission from one chronically infected individual (long branch in red). Figures **a** and **c** were created in BioRender.com.

To model the emergence and accumulation of mutations within each host, we use a standard, stochastic Wright-Fisher model ^34^. We assume that the pathogen population begins with a single starting genotype – consistent with tight transmission bottlenecks – and evolves in discrete generations subject to selection, mutation, and genetic drift. In each replication cycle, neutral and positively selected mutations are randomly introduced with rates *µ*_*N*_ and *µ*_*B*_, respectively. These mutation rates represent combinations of the basic probability per replication cycle that a new mutation is introduced and the probability that that mutation is beneficial or neutral. We assume that significantly deleterious mutations are rare enough to be efficiently eliminated by selection, and do not model them explicitly.

The distribution of fitness effects of beneficial and neutral mutations was derived from selection coefficients learned from SARS-CoV-2 temporal genomic data ^30^ (see **Supplementary Fig. 1**). In our model, the fitness effects of mutations are additive, so that the net increase or decrease in fitness *s*_*a*_ for a virus *a* with *n* mutations is

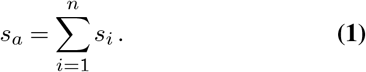

Here, the *s*_*i*_ are the selection coefficients that quantify the fitness effect of each mutation *i*. A positive selection coefficient indicates a beneficial mutation that increases fitness, while a negative coefficient indicates a deleterious mutation.

We assume that mutations that improve viral replication within a host also improve transmission between individuals. In principle, the effects of mutations on replicative fitness and transmission fitness can be different ^35^. As an example, some immune escape mutations generated during HIV-1 infection can be deleterious in other contexts, causing them to revert when the virus is transmitted to a new host ^36–40^. However, within-host mutations that produce fitness gains for replication and increase viral load can contribute to increased transmission, as has been shown for Spike mutations in SARS-CoV-2 (refs. ^41–43^). Furthermore, VOC mutations have been observed within individuals, including adaptive mutations concentrated in the Spike protein’s receptor binding domain and N-terminal domains ^25,26^. Despite these complications, we have aligned selection pressures within and between hosts for simplicity. Even in this simple case, complex evolutionary dynamics can occur.

We model transmission between individual donors and receptors of infection using a branching process that considers superspreading. In our model, the number of secondary infections is drawn from a negative binomial distribution *P*_*NB*_(*k, k/*(*k* + *R*^*i*^)), with *k* the dispersion parameter and 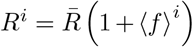 the effective reproductive number associated to the donor *i*. The negative binomial distribution has been used to model superspreading in past studies of viruses such as SARS and SARS-CoV-2 (refs. ^44–48^). Here, 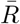 and ⟨*f* ⟩^*i*^ are the average baseline (reference) reproductive number and average fitness of the virus population from the donor host, respectively. As soon as infection is transmitted, donors are removed from the population. New infections are established by a single, randomly selected pathogen from the donor. Thus, most of the variant diversity previously generated is lost. This mimics the characteristic narrow transmission bottleneck observed in some pathogens, including SARS-CoV-2 (refs. ^7,8^).

The time between when an individual is first infected and when the infection is transmitted to a new host, which we refer to as the generation time, constrains the level of viral genetic diversity that can accumulate and be transmitted. The generation time varies based on the nature of the infection. Most infections are acute and cleared by the immune system in a short period, parameterized by *t*_*a*_. For SARS-CoV-2, we assume that two rounds of viral replication occur over each of the *t*_*a*_ days of infection ^49^. In immunocompromised hosts, infections can last far longer, with a generation time *t*_*c*_ ≫ *t*_*a*_. A longer generation time allows for more rounds of viral replication, facilitating the accumulation of genetic diversity. Each time an infection is transmitted, we take the probability that the new host develops a chronic infection to be *p*_*c*_ ≪ 1. The probability that a new infection is of short duration (acute) is then 1 − *p*_*c*_.

### Simulating pathogen evolution

We simulated multiple realizations of the evolutionary model over 1000 days. At each simulation time, we recorded the average number of mutations in transmitted variants (variants randomly sampled from within-host populations that are transmitted in new infections) and the number of chronically infected individuals across individual populations. Generation times for acute cases were set between 2 and 9 days, covering the values estimated from known infector-infectee transmission pair data or household data across different continents ^50,51^. For chronic cases, where generation times are less well determined, we sampled them from a log-normal distribution with mean *µ*_*L*_ = 150 days and standard deviation *σ*_*L*_ = 80 days. We maintained a fixed neutral mutation supply rate of *µ*_*N*_ = 10*−*4 mutations/cycle, based on the underlying mutation rate of SARS-CoV-2 (ref. ^49^) times the fraction of nonsynonymous mutations found neutral in the selection coefficient estimate from SARS-CoV-2 time series data ^30^ (Methods). We varied the beneficial mutation supply rate to explore its effect on pathogen evolutionary dynamics.

### Patterns of mutation accumulation

Within infected individuals, mutations accumulate progressively in viral populations over time (**Fig. 2**). Higher mutation rates naturally lead to a more rapid accumulation of mutations. Longer generation times (i.e., more generations of within-host evolution) also allow for more genetic diversity to accumulate within individuals, which can then potentially be transmitted to new hosts.

**Fig. 2.**
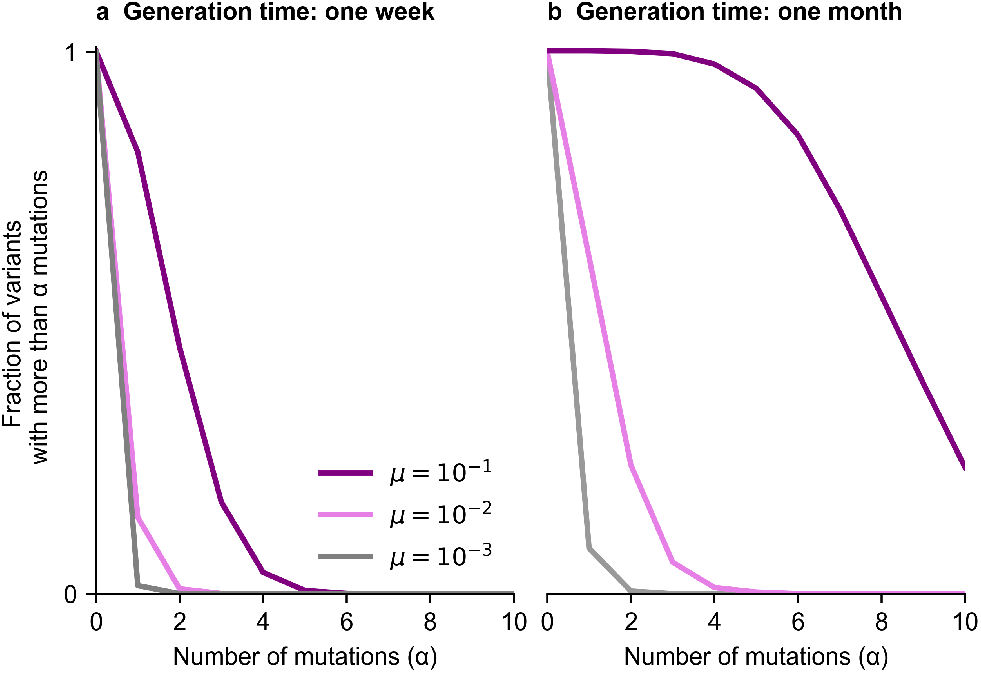
Genetic diversity after typical generation times of acute and chronic infections. **a**, Fraction of variants in the intra-host viral population that acquires more than *α* mutations over one week of infection. **b**, distribution of accumulated mutations after one month of infection. Higher mutation rates lead to the accumulation of more mutations. We consider the same rate, *µ*, for beneficial or neutral mutations.

Across infected individuals, we found that the evolution of viral populations fell into roughly three patterns (**Fig. 3**). In cases where there are few or no chronic infections, we observe few viral mutations (**Fig. 3a**). Single viral lineages tend to dominate the viral population with little competition between them.

**Fig. 3.**
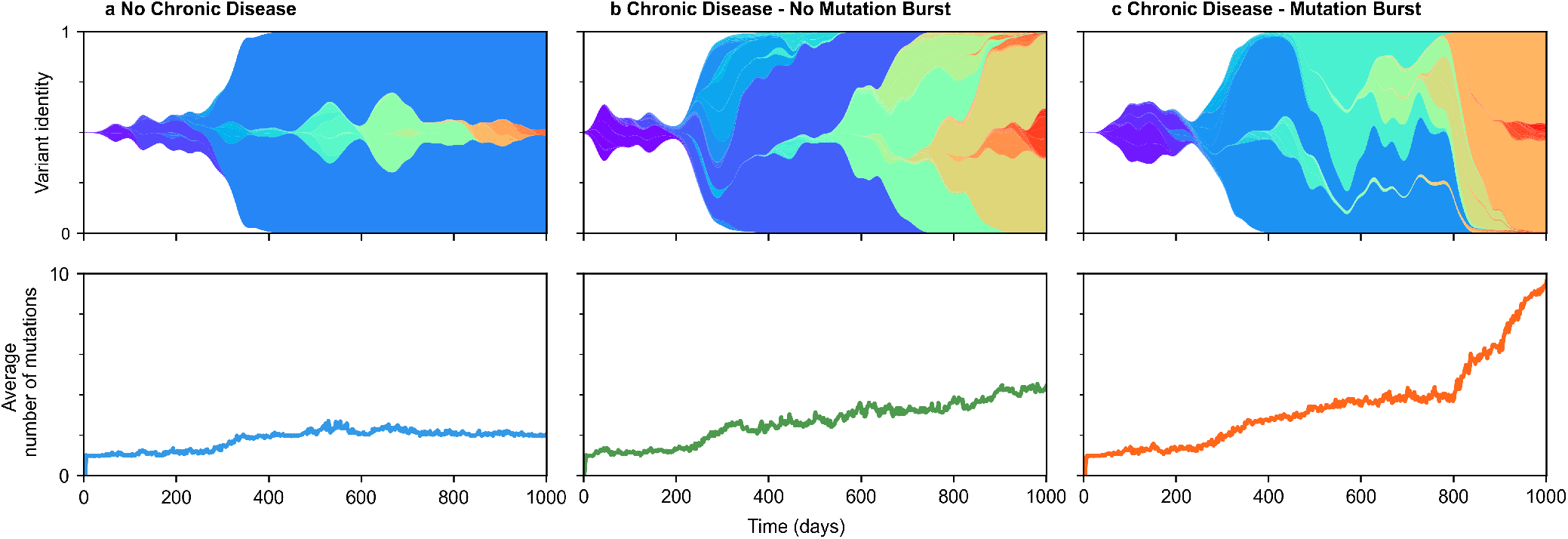
Evolution of viral variants under different scenarios. **a**, Dynamic evolution of variants within a population exclusively composed of acute cases. **b**, Population consisting of both acute and chronic cases but without mutation burst. **c**, Population consisting of both acute and chronic cases with one mutation burst at t=800 days. For all different simulations we consider beneficial and neutral mutation rates *µ*_*B*_ = 10^*−*3^ mutations/cycle and *µ*_*N*_ = 10^*−*4^ mutations/cycle respectively. For **b** and **c**, chronic cases are included in the percentage per transmission event *p*_*c*_ = 10^*−*3^ and generation times *t*_*c*_ are drawn from a log-normal distribution with mean *µ*_*L*_ = 150 days and standard deviation *σ*_*L*_ = 80 days.

When a significant number of chronic infections occur, two distinct outcomes are possible. In one case, the accumulation of mutations in viral populations accelerates and there is significant competition between viral lineages, but the increase in mutations over time remains roughly linear (**Fig. 3b**). In other simulations, we observe sudden “bursts” of mutations in viral populations, reminiscent of the emergence of SARSCoV-2 VOCs (**Fig. 3c**).

### Phase diagram for mutational bursts

To explore the relationship between parameter space and the emergence of mutational bursts, we generated a “phase diagram” of the number of mutational bursts per chronic disease case as a function of the model parameters (**Fig. 4**). To classify bursts versus linear accumulation of mutations, we first determined the distribution of maximum mutation accumulation rates across individuals in simulations without any chronic infections (Methods). We then identified bursts as events in which the rate of mutation accumulation was 3.5 or more standard deviations greater than the average maximum mutation accumulation rate in the simulations with only acute infections. We investigated a wide range of parameters, varying the fraction of chronic infections *p*_*c*_, acute generation times *t*_*a*_, and rate of beneficial mutations *µ*_*B*_ (see Methods).

**Fig. 4.**
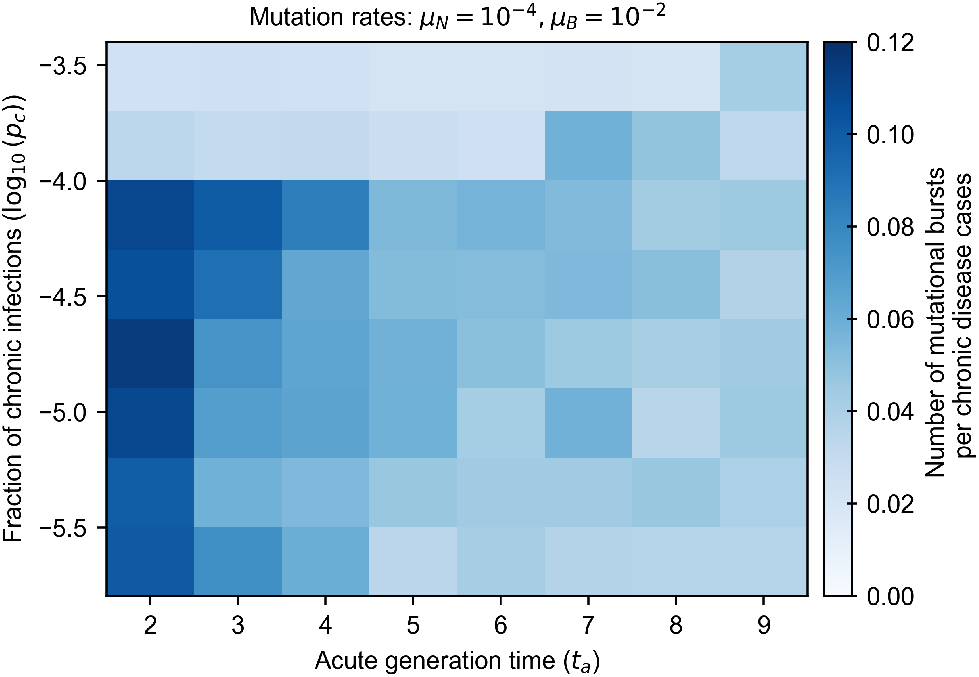
Number of mutational bursts per chronic disease case for beneficial mutation rate of 10^*−*2^ mutations/cycle. The frequency of mutational bursts decreases as acute generation times increase because chronic and acute subpopulations exhibit greater similarity, leading to homogenization within individual populations and reducing the likelihood of abrupt mutational events. The dependence on the fraction of chronic infections is nonlinear: while moderate levels of chronic infections lead to more frequent bursts, very high levels cause competition among multiple pathogen variants, each with numerous mutations, reducing the occurrence of isolated bursts. Each value represents an average over 1000 simulations.

For each choice of parameters, we computed the number of mutational bursts per chronic disease case over 1000 simulations.

We found several factors that facilitated the emergence of mutational bursts (**Fig. 4**). Intuitively, bursts occurred more frequently when beneficial mutation rates were higher (see analogous heatmap in **Supplementary Fig. 3** for a lower mutation rate). We also found that bursts occurred more frequently when the acute generation time *t*_*a*_ was shorter. Longer acute generation times lead to greater similarity in the viral populations in acute and chronically infected individuals, homogenizing the accumulation of mutations and decreasing the likelihood of abrupt increases in mutations.

Interestingly, we found that the likelihood of mutational bursts depends nonlinearly on the fraction of chronic infections. As the fraction of chronic infections increases, new adaptive mutations are generated more frequently and spread throughout the population, making isolated bursts unlikely. At very high frequencies of chronic infections, several pathogen variants with many mutations can be produced simultaneously. These variants then compete among hosts, reducing several potential bursts to a single one (see **Supplementary Fig. 4**).

### Effects of chronic infections on evolutionary rate

A recent study found that the evolutionary rate of SARSCoV-2 within a chronically infected individual was higher than the estimated global evolutionary rate of the virus, measured by the rate of substitutions over time ^25^. This can be attributed, in principle, to the absence of stringent bottlenecks imposed by transmission events. In our simulations, we observed that mutational bursts can occur due to the spread of new pathogen variants that evolved for long times within chronically infected individuals. Do chronic infections affect the overall evolutionary rate even in the absence of bursts?

To answer this question, we quantified the rate of accumulation of mutations across individuals over time in different scenarios (**Fig. 5**). Specifically, we measured the evolutionary rate within chronically infected individuals and the evolutionary rate between individuals in three different cases: in simulations with no chronic infections (*p*_*c*_ = 0), with chronic infections (*p*_*c*_ *>* 0) but without any observed mutational burst, and with chronic infections and at least one observed burst. Each measurement was averaged over 1000 simulations. Our results align with clinical data. Namely, the evolutionary rate within a single infected individual was higher than across the population of infected individuals in all cases.

**Fig. 5.**
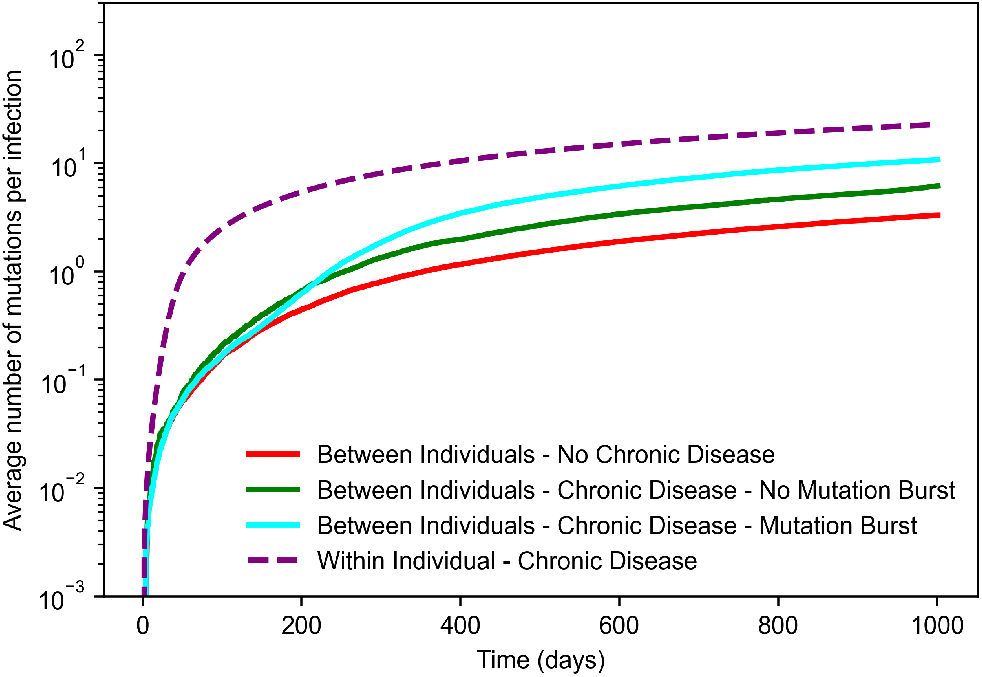
Average number of accumulated mutations per infection in simulations of within-host evolution and between-host evolution with and without chronic infections. For each observation time, the reported value represents the average number of mutations within intra-host viral populations of actively infected individuals, normalized by the total number of infected individuals at that time. The higher evolutionary rate is obtained within a chronically infected individual due to the absence of stringent bottlenecks imposed by transmission events. Between-host evolution with chronic infections, even in the absence of a burst, leads to an increased rate of mutation accumulation compared to populations with only acute infections. The simulations were conducted with a beneficial mutation rate of *µ*_*B*_ = 10^*−*3^ mutations/cycle, an acute generation time of *t*_*a*_ = 2 days, and a probability of new chronic infection of *p*_*c*_ = 4 *×* 10^*−*4^. Each curve represents an average of over 1000 simulations.

As expected, we found that the evolutionary rate between individuals was highest in populations with chronic infections and where at least one mutational burst was observed. However, even in the absence of a burst, the presence of chronic infections still leads to an increased rate of mutation accumulation compared to populations with only acute infections. Thus, chronic infections could still accelerate pathogen evolution through the generation and transmission of adaptive mutations, even without the production of SARS-CoV-2 VOC-like variants.

## Discussion

In this work, we modeled pathogen evolution within and between hosts including different types of infections: acute, short-term infections and rare chronic ones. The goal of our study was to understand how chronic infections can influence pathogen evolution over long times. Even with a simple, additive fitness landscape, we found that chronic infections can lead to SARS-CoV-2-like evolutionary dynamics, as mutants with multiple novel mutations arise and spread through the population. Such “bursts” of mutations were especially likely when acute generation times were short.

We found that the frequency of chronic infections had a strong and nonlinear effect on the frequency of mutational bursts. When chronic infections were rare, the number of observed mutational bursts scaled roughly linearly with the frequency of chronic infections. However, frequent chronic infections result in the generation and transmission of more adaptive mutations. Mutants with different beneficial mutations compete for hosts, making it more difficult for a single, dominant variant to quickly emerge.

Our model employs several simplifying assumptions that could be revisited in future work. First, we assumed that the fitness effects of pathogen mutations within and between hosts were the same. While certain mutations, such as those that increase viral load, are highly likely to improve both within-host replication and transmission between individuals, others may only be advantageous in particular scenarios. The distribution of fitness effects of mutations is also challenging to determine. Here, we used data from a recent study of SARS-CoV-2 evolution to parameterize our model ^30^. While the fitness effects of mutations in this study have extensive experimental support, they are subject to noise, and they were determined solely from between-host transmission rather than within-host replication. We have also assumed that the fitness effects of mutations are the same across hosts. Experiments ^52,53^ and computational analyses ^54^ have found many similarities between the fitness effects of mutations for genetically similar viruses, but some differences between hosts would be expected in real scenarios.

The model that we have developed is a type of “metapopulation” model ^55^, considering evolution both within and between hosts. Past studies have used such models to infer epidemiological dynamics ^56^ and explain phylogenetic structure ^57,58^, among other applications ^59^. Our work contributes to this area by exploring how different types of infections (i.e., acute versus chronic infection) contribute to pathogen evolutionary rates.

## ACKNOWLEDGEMENTS

The work reported in this publication was supported by the National Institute of General Medical Sciences of the National Institutes of Health under Award Numbers R35GM138233 and R35GM136381. A.R.D. acknowledges support from the National Science Foundation through the Physics Frontier Center for Living Systems (PHY-2317138).

## AUTHOR CONTRIBUTIONS

All authors contributed to methods development, data analysis, interpretation of results, and writing the paper. E.R.-H. and J.S. led computational analyses. A.R.D. and J.P.B. supervised the project.

## Methods

### Global evolutionary model

#### Within-host virus evolution model

The viral population, consisting of *N* infected cells, begins with a single starting genotype and undergoes discrete and non-overlapping Wright-Fisher generations subject to selection, mutation, and genetic drift. For each replication cycle, neutral and positive selected mutations are randomly introduced from a binomial distribution with rates {*µ*_*N*_, *µ*_*B*_} respectively. Once a mutation is generated for the genotype *a*, its selective effect *s*_*a*_ is drawn from distributions derived from experimental data. The genotype’s fitness is then updated as *f*_*a*_ = *f*^*wt*^ + *s*_*a*_, where *f*^*wt*^ represents the fitness of the wild-type genotype. Selected species in the subsequent generation are given by binomial sampling with success probability defined by their genotype fitness as *p*_*a*_ = *f*_*a*_*/ b f*_*b*_.

#### Between-host virus evolution model

In our model, new infections are established by one virion from the infection donor, and most of the variant diversity previously generated is lost. This mimics the characteristic narrow transmission bottleneck observed in respiratory viruses like SARS-CoV-2. The variant passed from the infection donor to an infected individual is randomly selected from the intrahost virus population. Subsequently, variants with higher fitness can persist through the transmission bottleneck only if there is sufficient intrahost evolution time to elevate their frequency. However, the most common scenario (with acute infections) is that variants without strong selective advantage overcome the transmission bottleneck by chance, a phenomenon of genetic drift.

Another relevant feature that our model incorporates is superspreading, where a small fraction of infectious hosts are responsible for most transmissions. For this, the number of secondary infections caused by an infected individual, at its generation period, is drawn from a negative binomial distribution *P*_*NB*_(*k, k/*(*k* + *R*^*i*^)), where *k* is the dispersion parameter and *R*^*i*^ = 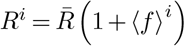 (1 + ⟨*f* ⟩^*i*^) represents the effective reproductive number associated to the host *i*. This is dependent on the average reproductive number 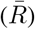 and the average fitness of the virus population from the donor host 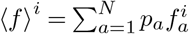 where *p*_*a*_ is the variant frequency. As soon as the infection occurs, donors are removed from the population, either due to death or immunity.

#### Dynamic simulations

We implemented the global evolutionary model in Julia. For the intra-host viral population size *N* = 1000 (ref. ^60^), we ran multiple independent realizations of the evolutionary model over 1000 days. The distribution of both neutral and beneficial mutation effects used during within-host replication cycles was fitted from the distribution of selection coefficients learned from SARS-CoV-2 temporal genomic data ^61^, as shown in **Supplementary Fig. 1**. To model generation times for chronic cases, we assume a log-normal distribution with mean *µ*_*L*_ = 150 days and standard deviation *σ*_*L*_ = 80 days. We select *k* = 1.0 as the dispersion parameter in the negative binomial distribution, which is within the estimated range for SARS-CoV-2^62^. We choose this moderate value to reflect that, in our model, transmission heterogeneity is not solely driven by *k*, as we consider an effective reproductive number *R*^*i*^ dependent on viral fitness in the donor, which also contributes to the non-homogeneous spread of secondary infections.

### Burst detection method

To identify bursts of mutations along the trajectories of accumulated mutations, we follow a step-by-step process. Initially, we calculate the slope at each time point for *M* trajectories obtained from simulations involving only acute cases. Subsequently, we apply a Savitzky–Golay filter ^63^ with a time window length (*w*) and polynomial order (*p*) to smooth the slope time series for each simulation. We extract the maximum values from the smoothed slope time series and use them to build a Gaussian null distribution (see **Supplementary Fig. 2a**).

We use the same smoothing process for the slope time series of simulations involving chronic disease cases. We then calculate the z-score for each time point using the mean and standard deviation obtained from the previously established null distribution. Mutation bursts are identified as outliers in this null distribution, defined by instances where the z-score exceeds 3.5. Given that multiple time points near the jump meet this criterion, we identify change points in the z-score time series. These change points delineate the start and end times of each jump, with the midpoint representing the burst time. The entire procedure is summarized in **Supplementary Fig. 2b**.

## Data and code

The data and code used in our analysis can be accessed from the GitHub repository at https://github.com/bartonlab/paper-SARS-CoV-2-evolution. This repository also contains the code used to analyze data and generate the figures presented in this paper.

---

**Supplementary Fig. 1.**
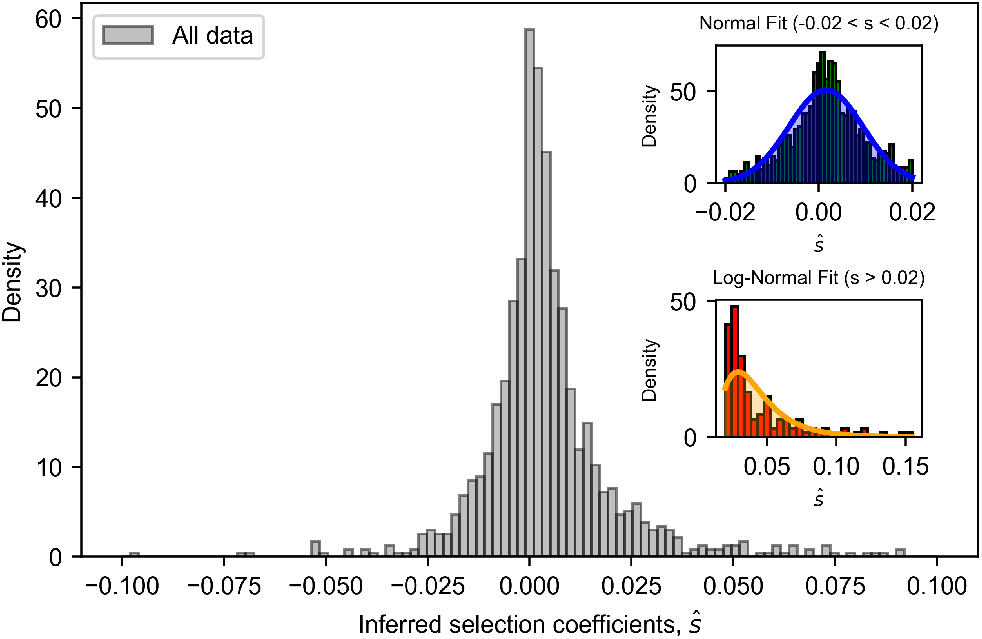
Inferred transmission effects of SARS-CoV-2 mutations. The main plot displays a histogram of selection coefficient values inferred from SARS-CoV-2 temporal genomic data ^61^. The top-right inset plot shows the normal distribution fit for coefficient values considered neutral (*−*0.02 *< ŝ<* 0.02); from this distribution, neutral mutation effects during within-host evolution were sampled. The bottom-right inset plot shows the log-normal distribution fit for values greater than 0.02, representing significantly beneficial mutations; from this distribution, beneficial mutation effects during within-host evolution were sampled.

**Supplementary Fig. 2.**
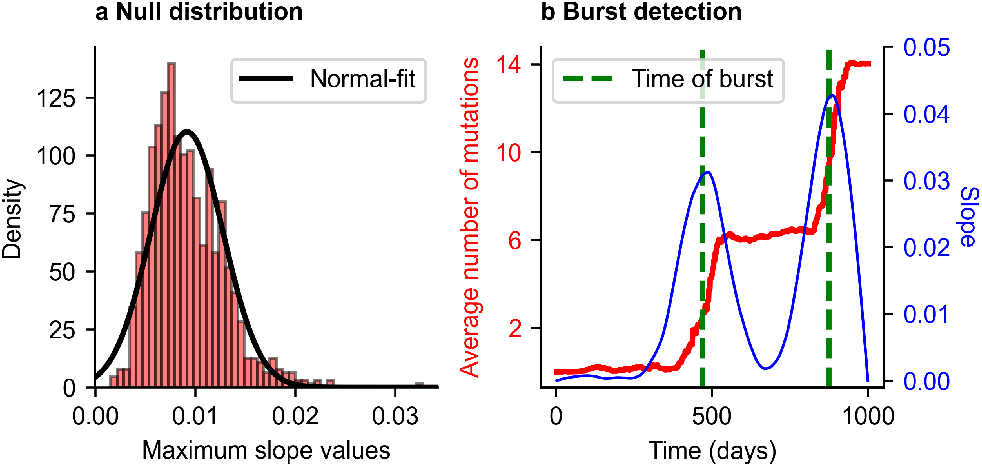
Detection of mutation bursts. **a**, Distribution of maximum slopes of accumulated mutation trajectories without chronic infection for acute generation time of *t*_*a*_ = 4.0 days and mutation rates: *µ*_*B*_ = 10^*−*3^ beneficial mutations per cycle and *µ*_*N*_ = 10^*−*4^ neutral mutations per cycle. **b**, For a simulation with chronic infection fraction *p*_*c*_ = 10^*−*4^, the number of accumulated mutations averaged over an individual’s population is shown in red. The blue curve indicates the smoothed slope time series with two peaks, detected by z-score time series change points and represented by the vertical green dashed lines. For smoothing using the Savitzky–Golay filter, we use parameters *w* = 150 and *p* = 1.

**Supplementary Fig. 3.**
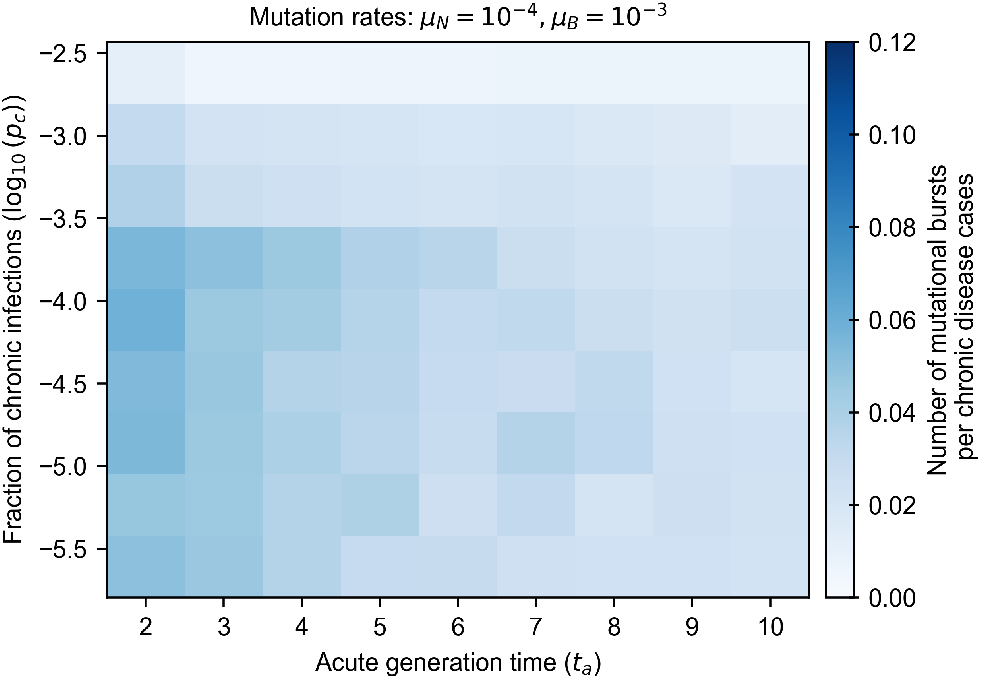
Number of mutational bursts per chronic disease case for beneficial mutation rate of 10^*−*3^ mutations/cycle. This figure is analogous to **Fig. 4** in the main text, but with a lower beneficial mutation rate.

**Supplementary Fig. 4.**
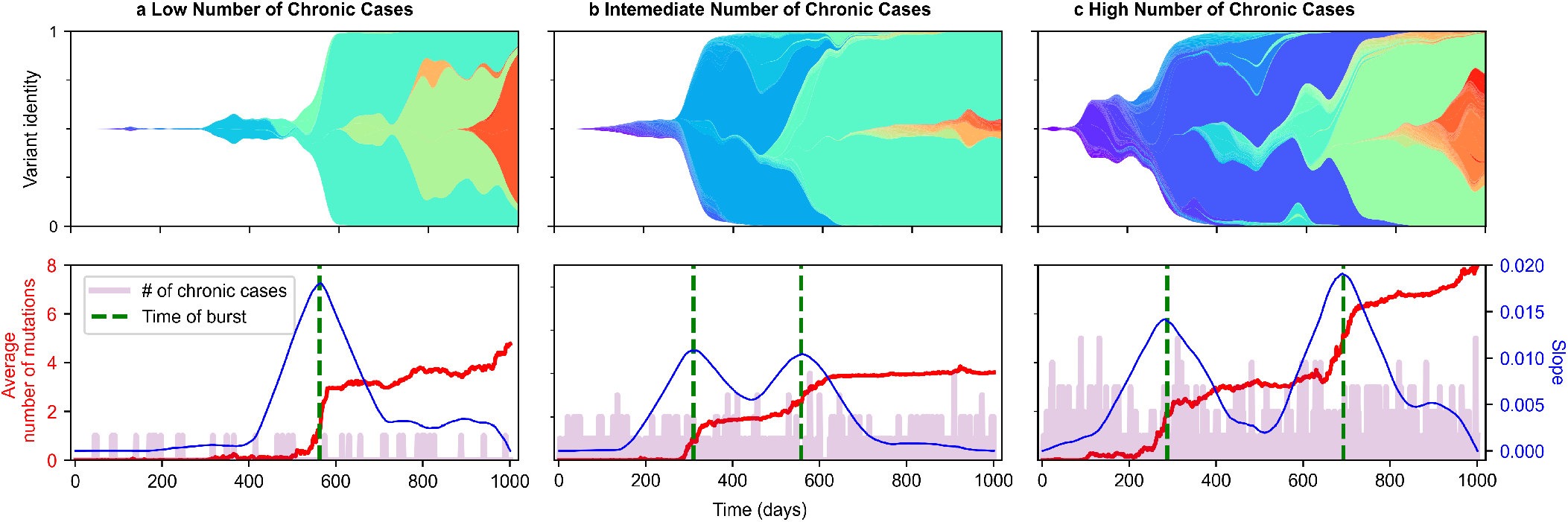
Dynamic evolution of the viral population under varying chronic infection probabilities. **a**, Low number of chronic cases, corresponding to a probability per transmission event of *p*_*c*_ = 4 *×* 10^*−*4^. **b** Intermediate number of chronic cases, with a probability per transmission event of *p*_*c*_ = 3.7 *×* 10^*−*3^. **c**, High number of chronic cases, resulting from a probability per transmission event of *p*_*c*_ = 7.0 *×* 10^*−*3^. For all simulations, we consider beneficial and neutral mutation rates *µ*_*B*_ = 10^*−*4^ mutations/cycle and *µ*_*N*_ = 10^*−*4^ mutations/cycle, respectively. Generation times are set at *t*_*a*_ = 2 for acute cases, while for chronic cases, they follow a log-normal distribution with a mean of *µ*_*L*_ = 150 days and a standard deviation of *σ*_*L*_ = 80 days.

